# Increased paternal corticosterone exposure preconception shifts offspring social behaviours and expression of urinary pheromones

**DOI:** 10.1101/2022.06.09.495572

**Authors:** Lucas B. Hoffmann, Evangeline A. McVicar, Rebekah V. Harris, Coralina Collar-Fernández, Michael B. Clark, Anthony J. Hannan, Terence Y. Pang

**Affiliations:** The Florey Institute of Neuroscience and Mental Health, Parkville, VIC, Australia; Florey Department of Neuroscience and Mental Health, Faculty of Medicine, Dentistry and Health Sciences, University of Melbourne, VIC, Australia; Department of Anatomy and Physiology, University of Melbourne, Parkville, VIC, Australia; Centre for Stem Cell Systems, Department of Anatomy and Physiology, University of Melbourne, Parkville, VIC, Australia

**Author notes:** Joint last authors (Anthony J. Hannan, Terence Y. Pang).

**Keywords:** paternal stress, epigenetic inheritance, social dominance, mate choice, major urinary protein, reproductive success

## Abstract

Studies have shown that paternal stress prior to conception can influence the innate behaviours of their offspring. The evolutionary impacts of such intergenerational effects are therefore of considerable interest. Our group previously showed that glucocorticoid treatment of adult male mouse breeders prior to conception leads to increased anxiety-related behaviours in male offspring. Here, we aimed to understand the transgenerational effects of paternal stress exposure on the social behaviour of progeny and its potential influence on reproductive success. We assessed social parameters including social reward, male attractiveness and social dominance, in the offspring (F_1_) and grand-offspring (F_2_). We report that paternal corticosterone-treatment was associated with increased display of subordination towards other male mice. Those mice were unexpectedly more attractive to female mice while expressing reduced levels of the key rodent pheromone Darcin, contrary to its conventional purpose. We investigated the epigenetic regulation of major urinary protein (*Mup*) expression by performing the first Oxford Nanopore direct methylation of sperm DNA in a mouse model of stress, but found no differences in *Mup* genes that could be attributed to corticosterone-treatment. Furthermore, no overt differences of the prefrontal cortex transcriptome were found in F_1_ offspring, implying that peripheral mechanisms are likely contributing to the phenotypic differences. Interestingly, no phenotypic differences were observed in the F_2_ grand-offspring. Overall, our findings highlight the potential of moderate paternal stress to affect intergenerational (mal)adaptive responses, informing future studies of adaptiveness in rodents, humans and other species.

## Background

Recent studies have demonstrated that the accumulation of paternal experiences before conception indirectly influence offspring behavioural phenotypes, and are largely attributed to epigenetic inheritance (1–5). This phenomenon has been studied in various animal models, with altered offspring phenotypes linked to paternal stress exposures persisting until the third generation of offspring (6). One common finding of studies to-date is a selective impact on offspring stress-relevant behaviours e.g. anxiety-like behaviour and social withdrawal. Studies of the potential mechanisms underlying these intergenerational effects have identified contributions of distinct subpopulations of non-coding RNAs. For example, through microinjection studies of fertilised oocytes and zygotes, paternal preconception stress-associated changes to sperm small non-coding RNAs in mice were found to influence anxiety-related behaviours and the stress-induced corticosterone response of adult F_1_ offspring (7,8). There is also emerging evidence of a specific contribution of sperm long non-coding RNAs to influence adult F_1_ offspring metabolic phenotypes (9). Thus, paternal stress-driven intergenerational adaptations could be a contributor to the ‘missing heritability’ problem associated with anxiety, depression and other psychiatric disorders in humans (10). Additionally, some speculate about the evolutionary advantages that such heritability could confer, such as adding phenotypic variation (11,12).

Our previous work on the paternal corticosterone-supplementation model of generalised daily stress had reported elevated anxiety-like behaviours of male F_1_ offspring (PatCort) and the emergence of depressive-like behaviours in male F_2_ grand-offspring (GPCort) (13). We subsequently found that PatCort mice were resistant to the anxiolytic effects of environmental enrichment (routinely reported in the wider literature) and had reduced sensitivity to the selective serotonin reuptake inhibitor sertraline (14). Other independent preclinical studies of distinct mouse models of stress have also found defects in sociality and social recognition accompanying impaired serotonergic signalling (15), as well as dysregulation of the physiological stress response (8). In rodents, appropriate social behaviour is particularly important for reproduction and survival, and thus influences individual fitness. Given the increasing evidence that epigenetic inheritance influences behavioural endophenotypes, it is possible that epigenetic inheritance also underlies social behaviours relevant to successful reproduction, with consequences for adaptivity and species evolution (16,17).

Here, we embarked on a transgenerational study of rodent social behaviours highly relevant to their reproductive success. We investigated social dominance and male attractiveness across two generations of progeny in the paternal corticosterone-supplementation model of paternal stress. We hypothesised that a paternal history of stress would result in offspring displaying lower preference for social reward, increased subordinate behaviour during social interaction, and reduced preference from potential female mates. We followed up on these findings by investigating the expression of major urinary proteins (MUPs), which are non-volatile pheromones (18). Here, we present evidence that paternal stress exposure preconception can exert an intergenerational influence over behavioural endophenotypes that determine reproductive success in offspring. Our initial molecular and epigenetic studies also highlight the complex involvement and regulation of major urinary proteins (MUPs) in rodent social behaviour, and the regulation of MUP expression by the paternal exposure to corticosterone.

## Results

### Corticosterone (Cort)-treatment diminishes adult male social dominance but does not adversely affect mate attraction

We first evaluated the effect of Cort-treatment on the social behaviours of adult male mice. 4 weeks of Cort-treatment significantly reduced the percentage of wins in the tube test (Figure 1A, χ*^2^*=30.25, *p*<0.001), which did not correlate to their body weight (Supplementary Figure S3). Based on this, we expected that Cort-treated males would be less preferred by potential female mates. However, there were no differences across all three variations of the test we conducted (‘Standard’ set up: Sum of Signed Ranks *W*=8.000, *p*=0.8603, Figure 1B; ‘Mouse only’ set up: Sum of Signed Ranks *W*=32.00, *p*=0.4332, Figure 1C; ‘Bedding only’ set up: Sum of Signed Ranks *W*=36.00, *p*=0.3755, Figure 1D). Based on interaction times, females did not show preference for one group of males over the other (Figure 1E, ‘Standard’ set up: χ*^2^*=0.2500, *p*=0.6171; Figure 1F, ‘Mouse only’ set up: χ*^2^*= 1.000, *p*=0.3173; Figure 1G, ‘Bedding only’ set up: χ*^2^*=0.2500, *p*=0.6171). Therefore, we found that Cort-treatment reduces social dominance and does not affect male attractiveness.

**Figure 1.**
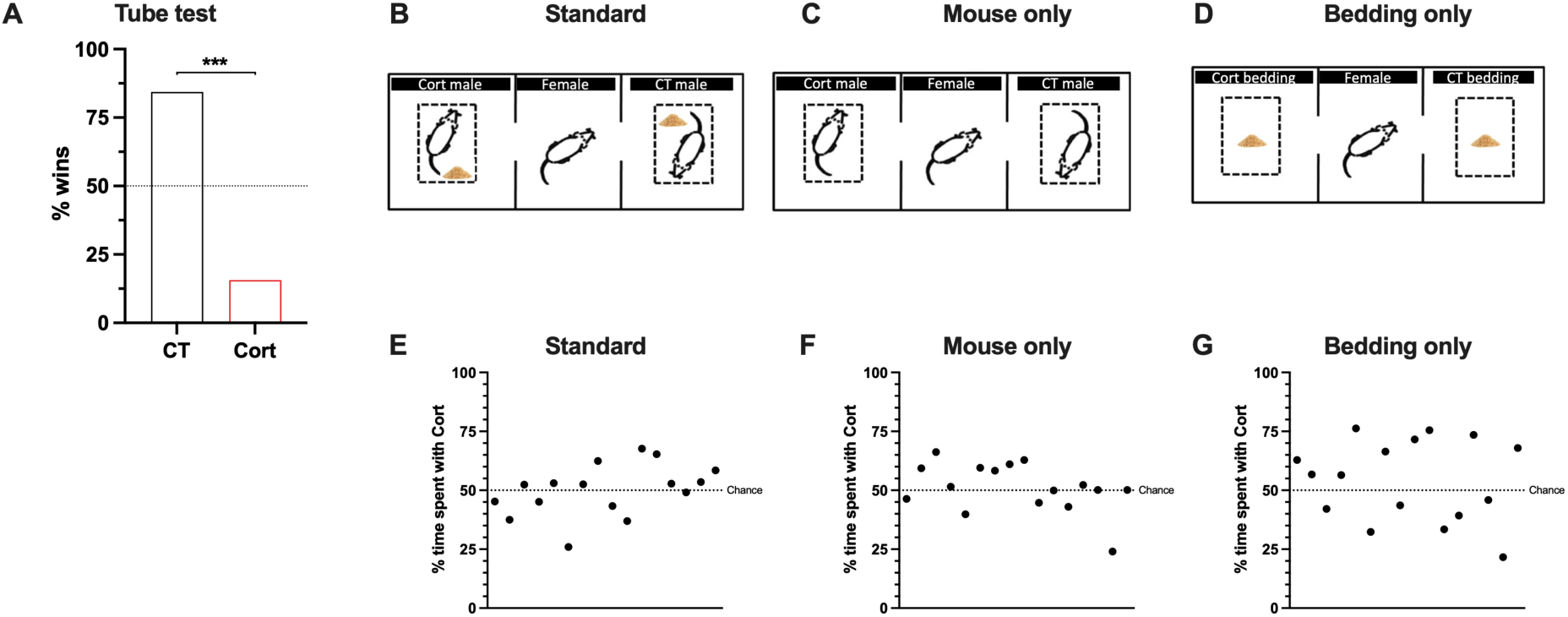
Assessing effects of Cort-treatment on social dominance and mate-choice attractiveness. (A), corticosterone treatment reduces male dominance as assessed by the percentage of wins in the tube test. % wins calculated as the number of wins per group in the total number of interactions. Total of 64 unique interactions. (B – D), figures adapted from Toth and Neumann, 2013. In all of them, a female mouse is represented in the centre of a 2-chamber apparatus. In figure B, male mice from CT and Cort group are located at each end of the apparatus, alongside soiled bedding from their home cage. In figure C, male mice from CT and Cort group are presented without bedding. In figure D, only soiled bedding from CT and Cort cages is presented. (E – G), male attractiveness was not affected by corticosterone-treatment. Each point represents the results from one female mouse. A, E – G: one-sample chi-squared test. *** *p*<0.001.

### Paternal Cort-treatment affects social dominance and mate attraction of adult male offspring

Juvenile (PND 35) F_1_ male and female PatCort offspring displayed a clear preference for the social-conditioned bedding (Figure 2A, Males: *F*(*_1,72_*)=12.76, *p*<0.001; Figure 2C, Females: *F*(*_1,74_*)=8.227, *p*=0.0054). No effects of paternal treatment were found for males (Figure 2B, *U*=601, *p*=0.3811) nor for females (Figure 2D, *U*=640.5, *p*=0.4035). Therefore, F_1_ PatCort offspring mice displayed normal preference for social reward, and it was independent of the paternal corticosterone exposure.

**Figure 2.**
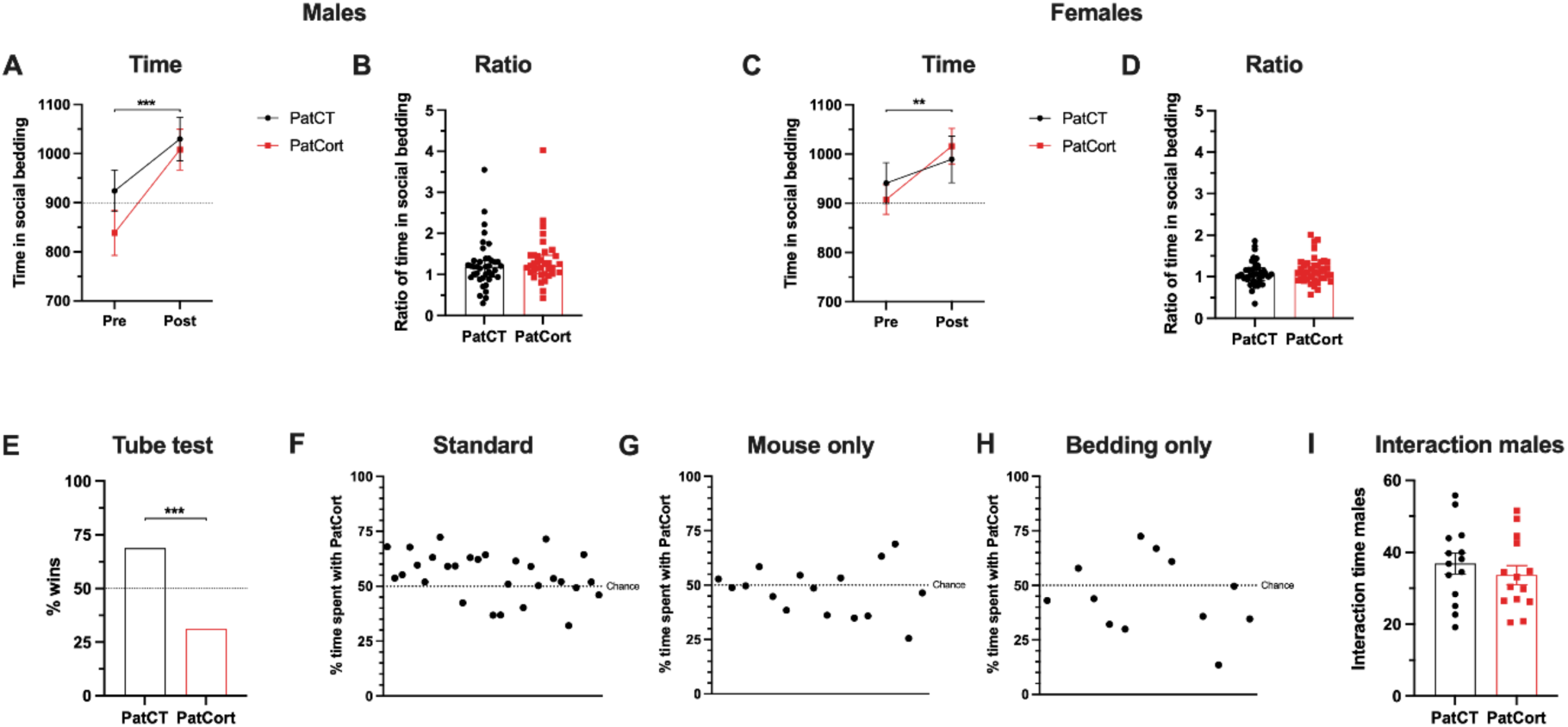
Assessing effects of paternal Cort-treatment on offspring social behaviour. (A – D), the degree of social reward in male or female offspring was not affected by paternal Cort-treatment. Time and ratio of the time spent in the social-conditioned bedding. (E), paternal Cort-treatment reduces male dominance in the tube test. % wins calculated as the number of wins per group in the total number of interactions. Total of 112 unique interactions. (F – H), paternal corticosterone treatment increases male attractiveness in the ‘standard’ set up only. Each point represents the results from one female mouse. (I), PatCort mice do not interact more with female mice, compared to PatCT. A and C: 2-way ANOVA, data represented as mean ± SEM. B and D: Mann-Whitney test, data represented as median ± interquartile range. E – H: one-sample chi-square test. I: unpaired t-test, data represented as mean ± SEM. ** *p*<0.01, *** *p*<0.001.

In the tube test, adult F_1_ male PatCort offspring recorded significantly fewer winning interactions, indicative of a lower order of social dominance (Figure 2E, χ*^2^*=15.75, *p*<0.001). In the mate-choice test, *naïve* females investigated F_1_ male PatCort offspring for significantly longer periods than their PatCT counterparts (Sum of Signed Ranks *W*=241.0, *p*=0.0080), and a significantly higher percentage of time (Figure 2F, χ*^2^*=7.759, *p*=0.0053). No differences were observed between the groups for the ‘mouse only’ nor the ‘bedding only’ set ups (‘Mouse only’: Sum of Signed Ranks *W*=-40.00, *p*=0.3225; ‘Bedding only’: Sum of Signed Ranks *W*=-18.00, *p*=0.5186), and the total percentage of time was comparatively similar between the groups (Figure 2G, ‘mouse only’ set up, χ*^2^*=1.000, *p*=0.3173; Figure 2H, ‘bedding only’ set up, χ*^2^*=1.333, *p*=0.2482). Additionally, analysis of the total time each male directly interacted with the female when she approached them revealed no differences (Figure 2I, *t*(*_26_*)=0.8106, *p*=0.4249). Thus, paternal Cort-treatment was associated with intergenerational shifts in social-relevant behaviours of adult F_1_ male PatCort offspring.

### Paternal Cort-treatment effects do not transgenerationally alter grand-offspring social behaviours

Juvenile F_2_ male and female grand-offspring showed a preference for the social-conditioned bedding (Figure 3A, Males: *F*(*_1,76_*)=35.14, *p*<0.001; Figure 3C, Females: *F*(*_1,80_*)=12.09, *p*<0.001). No differences in social reward were found between the groups for males (Figure 3B, *U*=714.5, *p*=0.6527) nor for females (Figure 3D, *U*=755, *p*=0.4547).

**Figure 3.**
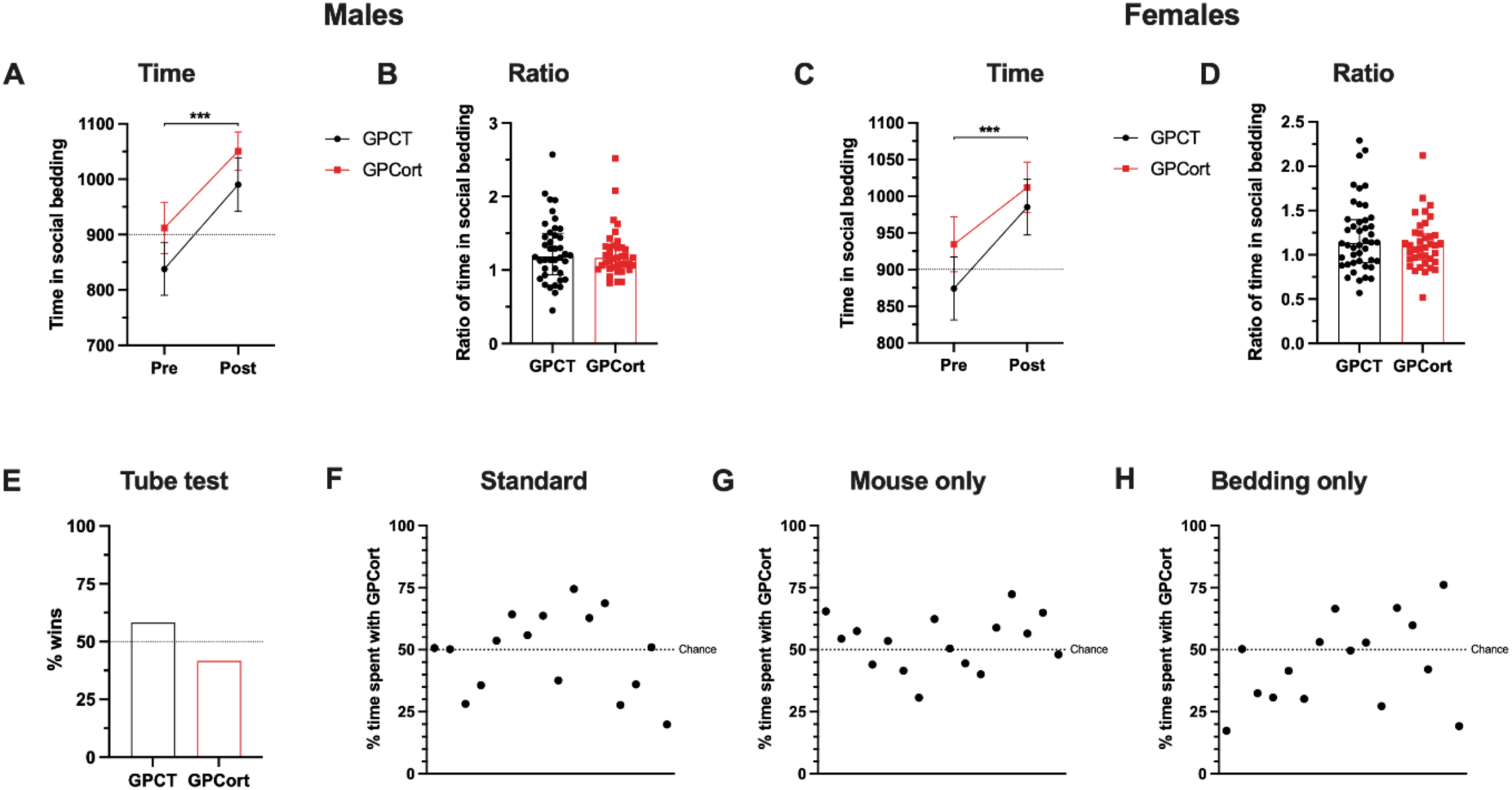
Assessing effects of grand-paternal Cort-treatment on grand-offspring social behaviour. (A – D), the degree of social reward in male or female grand-offspring was not affected by grand-paternal Cort-treatment. Time and ratio of the time spent in the social-conditioned bedding. (E), social dominance in male grand-offspring was not affected by grand-paternal Cort-treatment. % wins calculated as the number of wins per group in the total number of interactions. Total of 64 unique interactions. (F – H), male attractiveness in male grand-offspring was not affected by grand-paternal Cort-treatment. A and C: 2-way ANOVA, data represented as mean ± SEM. B and D: Mann-Whitney test, data represented as median ± interquartile range. E – H: one-sample chi-squared test. *** *p*<0.001.

In the tube test, GPatCort and GpatCT groups recorded similar numbers of wins (Figure 3E, χ*^2^*=2.667, *p*=0.1025). In the mate-choice test, females spent similar amounts of time interacting with both groups of mice across all of the set ups (Sum of Signed Ranks *W*=-10.00, *p*=0.8209; ‘Mouse only’ set up: Sum of Signed Ranks *W*=40.00, *p*=0.3160; ‘Bedding only’ set up: Sum of Signed Ranks *W*=-40.00, *p*= 0.3225). The percentage time spent by the female interacting with both groups was similar (Figure 3F, ‘Standard’ set up: χ^2^= 1.000, *p*= 0.3173; Figure 3G, ‘Mouse only’ set up: χ^2^= 1.000, *p*= 0.3173; Figure 3H, ‘Bedding only’ set up: χ^2^= 0.2500, *p*= 0.6171). Therefore, no transgenerational effects on social-relevant behaviours of F_2_ GPCort grand-offspring were observed in association with paternal Cort-treatment.

### Paternal Cort-treatment alters the expression of a subpopulation of MUPs in F_1_ offspring

Since pheromones play an important role in rodent male social hierarchy and mate attraction, we quantified the expression profiles of MUPs in the urine of F_1_ male offspring.

We first assessed urinary creatinine content to account for urinary dilution, and determined that it was not different between PatCT and PatCort male offspring (Figure 4A, *t_(12.74)_*=1.704, *p*=0.1126), which indicates that their urination volume did not differ (19). The relative levels of total MUPs also did not differ between the groups (Figure 4B, *t_(15.24)_*=1.792, *p*=0.0930). Next, we differentiated between the three major MUP bands (Figure 4C), in agreement with previous publications (18,20). Semi-quantification of the protein bands revealed that PatCort males had reduced concentrations of the small MUP band (Figure 4D, *t_(15.41)_*=2.252, *p*=0.0393) and of Darcin (Figure 4F, *U*=53, *p*=0.0243). No difference in the big MUP band was observed (Figure 4E, *t_(14.09)_*=1.904, *p*=0.0775). Additionally, the variability of all three bands of MUP proteins was higher in the PatCort mice (Small MUP band: *F*(*_12,15_*)=5.693, *p*=0.0022; Big MUP band: *F*(*_12,15_*)=9.406, *p*=0.0001; Darcin: *F*(*_12,15_*)=5.614, *p*=0.0024; F test to compare variances). Therefore, there appears to be some selectivity in terms of the intergenerational impacts of paternal Cort-treatment on offspring production of MUPs.

**Figure 4.**
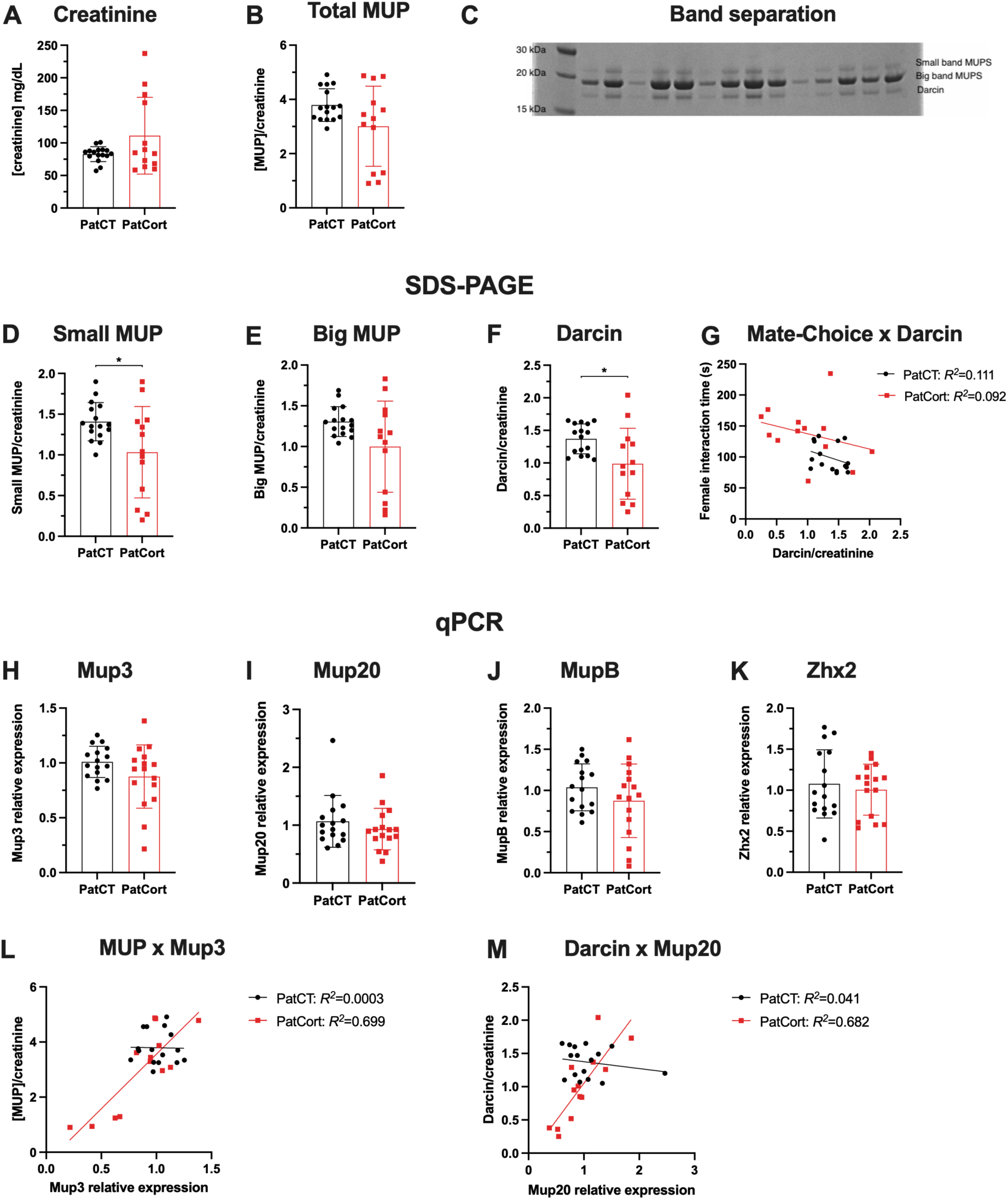
Assessing effects of paternal Cort-treatment on offspring urinary MUP levels and liver gene expression. (A – B), urinary creatinine or total MUP were not affected by paternal Cort-treatment. (C), after SDS-PAGE of mouse urine, three different MUP bands can be seen, whose molecular weight correspond to previously published literature. Heavier band = ‘Small MUP’, middle band = ‘Big MUP’, lighter band = ‘Darcin’. (D – F), paternal Cort-treatment induces reduced specific MUP populations in the urine. (G), time being investigated by the female does not correlate with urinary Darcin concentration. (H – K), *Mup* gene expression in the liver was not affected by paternal Cort-treatment. (L – M), *Mup* genes correlate with specific MUP populations in PatCort, but not in PatCT. A – B: unpaired t-test with Welch correction. D and E: unpaired t-test with Welch correction. F: Mann-Whitney test. G: correlation of Pearson and simple linear regression. H: unpaired t-test with Welch correction. I: Mann-Whitney test. J and K: unpaired t-test. L – M: correlation of Pearson and simple linear regression. Data represented as mean ± SD. * *p*<0.05.

Darcin was proposed to be the key MUP subtype to act as a female attractant pheromone (18). However, urinary Darcin concentrations of male mice of both PatCT and PatCort groups were not significantly correlated with total time being investigated by the female in the mate-choice test (Figure 4G, PatCT: Pearson’s correlation *R^2^*=0.1112, *p*=0.2068; PatCort: Pearson’s correlation *R^2^*=0.0921, *p*=0.3134).

Since MUP proteins are produced and secreted by the liver (21), we further investigated hepatic gene expression of various *Mup* genes and *Zhx2* (a transcript factor for *Mup* genes) (22). However, *Mup* gene expression in PatCort mice did not differ from PatCT mice (Figure 4H, *Mup3*: *t_(21.88)_*=1.666, *p*=0.1100; Figure 4I, *Mup20*: *U*=107, *p*=0.4393; Figure 4J, *MupB*: *t*(*_30_*)=1.235, *p*=0.2264), despite the increased variability in the gene expression of *Mup3* (*Mup3*: *F*(*_15,15_*)=4.117, *p*=0.0094; *Mup20*: *F*(*_15,15_*)=1.555, *p*=0.4020; *MupB*: *F*(*_15,15_*)=2.444, *p*=0.0939; F test to compare variances) in PatCort mice. There was also no difference in hepatic *Zhx2* gene expression (Figure 4K, *Zhx2*: *t*(*_30_*)=0.5436, *p*=0.5907).

*Mup* gene expression levels displayed a strong positive correlation with MUP protein populations in the PatCort offspring. *Mup3* correlates very strongly with total MUP urinary concentration in the urine of PatCort mice (Figure 4L, Pearson’s correlation *R^2^*=0.6988, *p*<0.001). Interestingly, this correlation was not observed for PatCT mice (Figure 4L, Pearson’s correlation *R^2^*= 0.0003, *p*=0.9498). *Mup20*, in its turn, correlates very strongly with the Darcin band population of MUPs in the urine of PatCort mice (Figure 4M, Pearson’s correlation *R^2^*=0.6823, *p*=0.0005), but did not correlate in PatCT mice (Figure 4M, Pearson’s correlation *R^2^*=0.041, p=0.4521). The Darcin band has been attributed to the *Mup20* gene in previous publications (18,19). These findings suggest that in PatCort mice, the high variability in the gene expression of *Mup3* is likely causing the high variability in the protein level.

### MUP profile is not transgenerationally influenced by paternal Cort-treatment

We also assessed urinary creatinine and MUP protein concentrations of F_2_ male grand-offspring mice. Both urinary creatinine (Figure 5A, *U*=101.5, *p*=0.8897) and total MUP concentrations (Figure 5B, *t*(*_27_*)=0.9832, *p*=0.3343) did not significantly differ between the groups. No between-group differences were also detected for any of the three major MUP bands, namely small MUPs (Figure 5C, *t*(*_27_*)=0.4547, *p*=0.6529), big MUPs (Figure 5D, *t*(_27_)=0.7838, *p*=0.4400) and Darcin (Figure 5E, *t*(*_27_*)=0.5951, *p*=0.5568). Additionally, there were no differences in variability between the groups.

**Figure 5.**
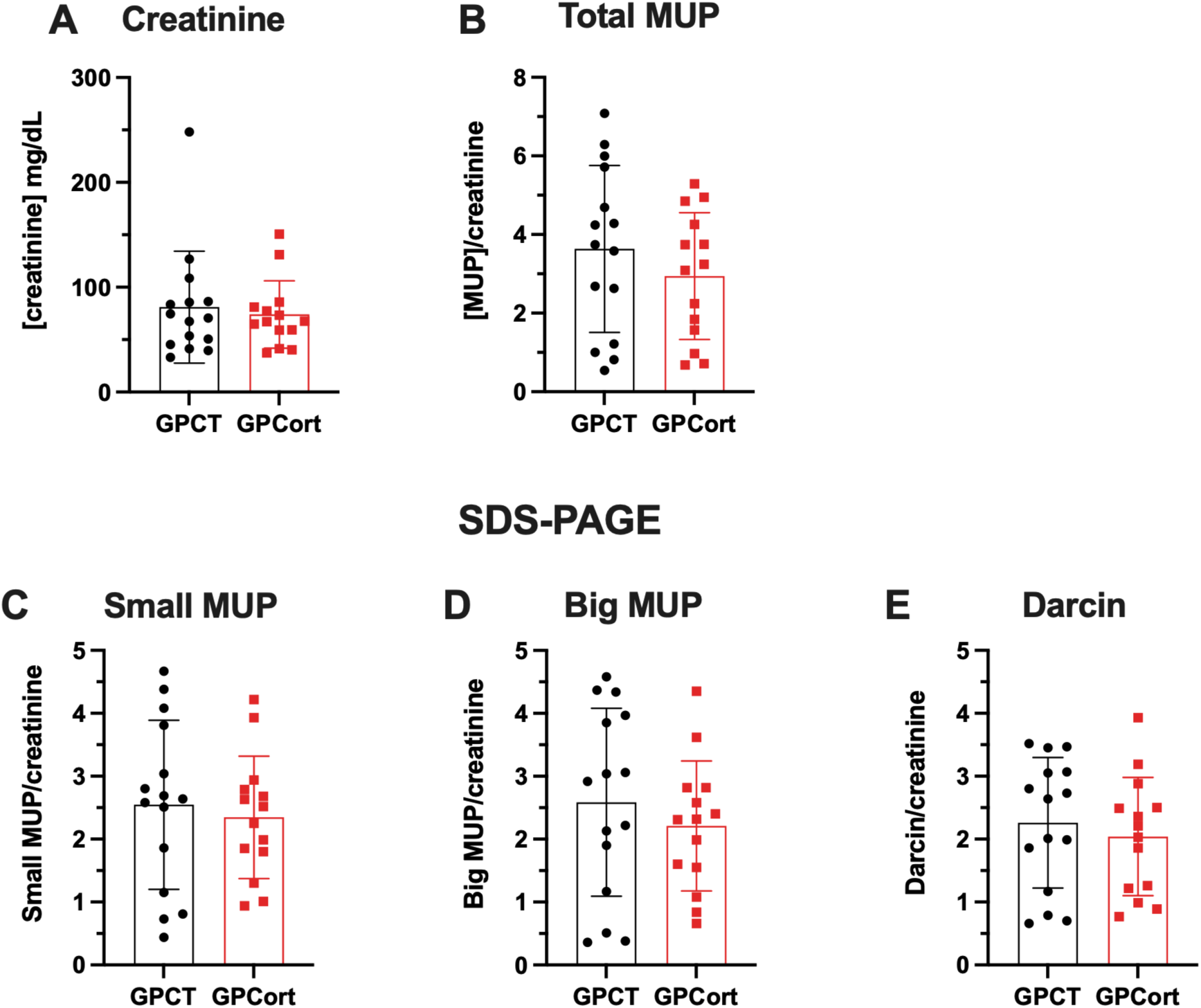
Assessing effects of grand-paternal Cort-treatment on grand-offspring urinary MUP levels. (A – B), urinary creatinine or total MUP were not affected by grand-paternal Cort-treatment. (C – E), specific MUP populations in the urine were not affected by grand-paternal Cort-treatment. A: Mann-Whitney test. B – E: unpaired t-test. Data represented as mean ± SD.

### Cort-treatment does not alter *Mup* gene DNA methylation

Differential methylation of *Mup* genes has been reported, but the epigenetic regulation of *Mup* gene expression remains unclear (23–25). To address this, we developed an optomised sperm DNA extraction protocol that enabled us to conduct the first long-read nanopore sequencing study of *Mup* gene methylation in sperm DNA harvested from Cort-treated and control F_0_ males. Overall, CpG methylation indicated a very high methylation frequency across the genome (not shown), which is expected for sperm since it is a transcriptionally quiescent cell population. CpG methylation located within the *Mup* gene cluster was also found to be highly methylated (Figure 6H), with no obvious differences between both groups (Figure 6A). We noted a potential pattern of increased methylation associated with Cort-treatment at the promoter region of *Mup20* and decreased methylation in the 3’ downstream region of the gene (Figure 6B); these would be consistent with decreased gene expression of *Mup20* and decreased expression of Darcin protein we had found (26). Therefore, we assessed CpG methylation at the promoters of the *Mup20* gene, whose locations were obtained from the UCSC genome browser promoter track. However, no differences in methylation were found (*p*=0.96 – Figure 6J). We further inspected methylation of the transcription factor *Zhx2* and additional *Mup* genes of interest (*Mup3*, *Mup2*, *Mup15*, *Mup18* – Figures 6C – 6G,) identifying no major differences. Consistent with this, an analysis of differentially methylated regions (DMRs) using DSS revealed no significant differences between the groups at the *Mup* locus. Thus, it appears that DNA methylation is not a major epigenetic regulator of *Mup* expression in this rodent model.

**Figure 6.**
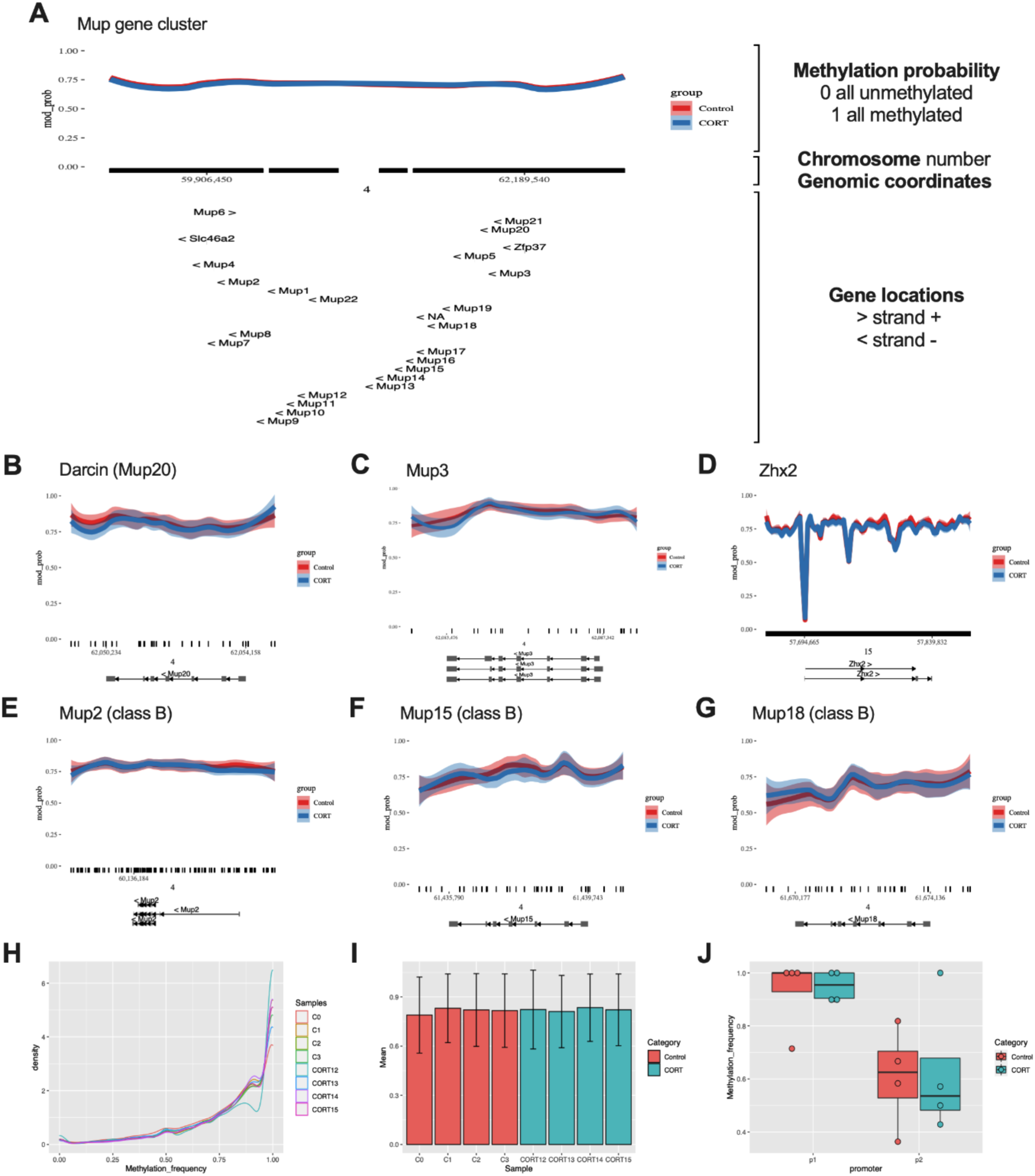
DNA methylation profile of *Mup* genes. (A), whole *Mup* cluster genomic region. (B), *Mup20*. (C), *Mup3*. (D), *Zhx2*. (E), *Mup2*. (F), *Mup15*. (G), *Mup18*. (H), *Mup* cluster methylation density plot. (I), *Mup20* gene methylation frequency per sample. (J), *Mup20* promoters 1 and 2 methylation frequency. Methylation frequency=1: Methylated CpG. Methylation frequency=0: Unmethylated CpG.

### Adult male offspring prefrontal cortex gene expression is relatively unchanged by paternal Cort-treatment

The rodent prefrontal cortex is heavily implicated in displays of social dominance (27), as well as anxiety-relevant behaviours that we have reported in this model (13,28). We therefore conducted transcriptome profiling of this brain region to determine whether gene expression differences underlie the F_1_ offspring phenotypes we observed. Overall, we found that samples were very similar, independent of their group (Figure 7A). Only 32 genes were found to have *p* < 0.05 and log-fold change (LFC) threshold = 1, with 3 upregulated and 29 downregulated in the PatCort male mice (Figure 7B, Table 2), although they were not statistically significant after FDR correction. Similarly, Gene Set Enrichment Analysis (GSEA) was performed, but no significant gene sets were found (Table 3).

**Figure 7.**
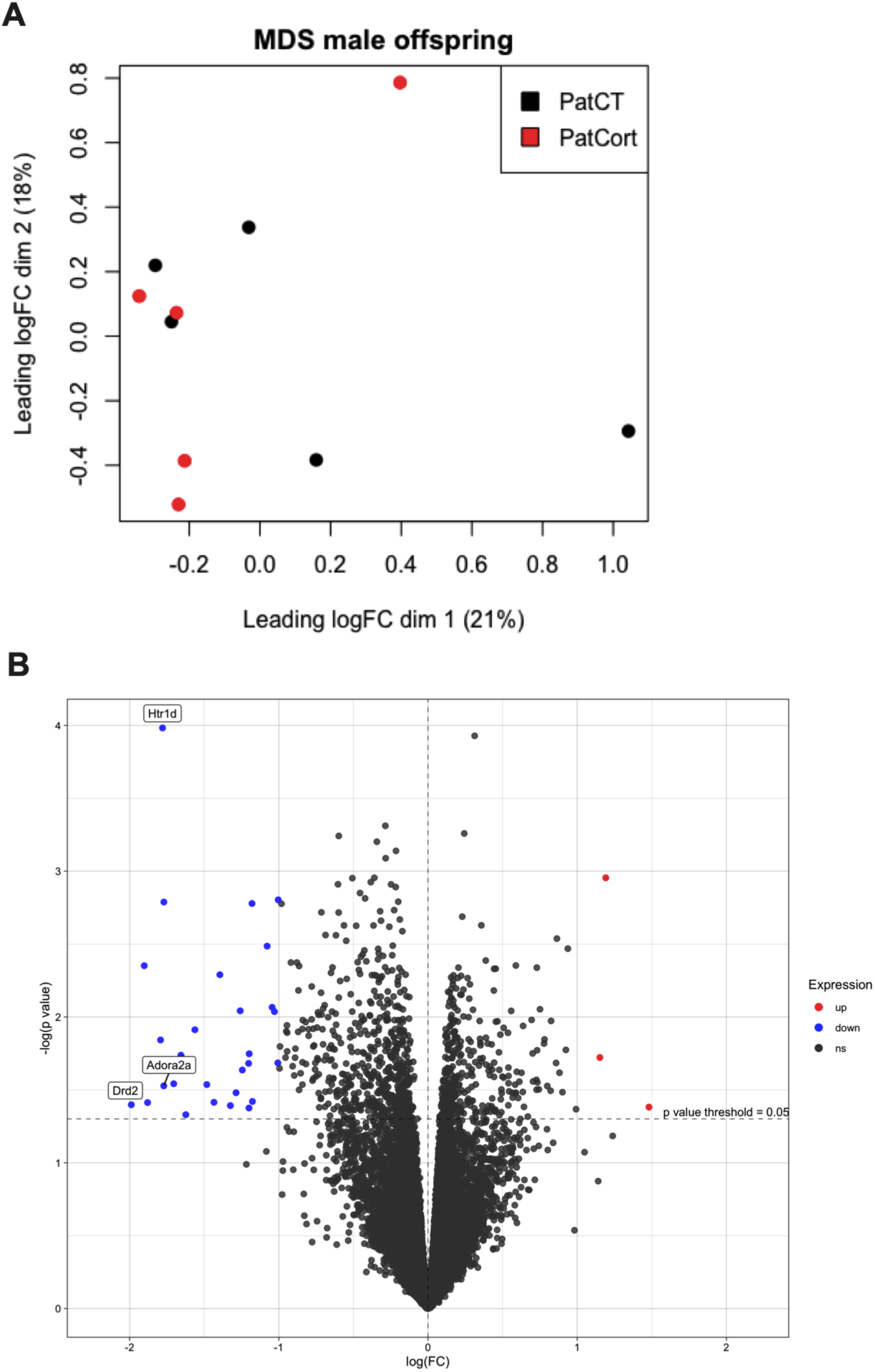
Assessing effects of paternal Cort-treatment on offspring prefrontal cortex gene expression. (A), multidimensional scaling plot of distances between the male F_1_ offspring prefrontal cortex gene expression profiles. (B), volcano plot showing genes with *p*<0.05 and log-fold change threshold of 1 prior to FDR correction. Red: genes upregulated in PatCort. Blue: genes downregulated in PatCort, compared to PatCT.

## Discussion

This study has uncovered novel evidence of the paternal influence over offspring social behaviours. Paternal Cort-treatment was associated with reduced social dominance and increased attractiveness of their male F_1_ offspring, in addition to the elevated anxiety phenotype we previously reported (13). Interestingly, these effects on social behaviours were restricted to the immediate generation (F_1_ offspring), with no significant transgenerational effects on the F_2_ grand-offspring. Additionally, the male F_1_ PatCort offspring also showed reduced and more variable MUP protein output in their urine, in particular the band that corresponds to the male pheromone Darcin. By performing the first sperm DNA methylome sequencing study in mice treated with corticosterone, we established that the abnormal MUP expression is not a result of dysregulated DNA methylation of the various *Mup* genes. Additionally, the absence of differences in the PFC transcriptome strongly suggests that the altered social responses of F_1_ PatCort offspring likely originate in other brain regions key to social interaction, such as the anterior cingulate cortex or the ventromedial hypothalamus nucleus (29,30).

### F_0_ male mice showed reduced dominance and no difference in attractiveness

Male mice treated with corticosterone in their drinking water for 4 weeks showed a large reduction in their social dominance, with no effects on their attractiveness to female mice. There is extensive literature on dominance and social hierarchy priming response to stress (31), as it is known that hierarchical rank modulates the hypothalamic-pituitary-adrenal (HPA) system reactivity (32), possibly due to higher-order impacts of the prefrontal cortex on hypothalamic function. Studies on stressful exposures modulating social dominance and hierarchical rank are less studied, but it has been reported that a cortico-hypothalamic circuit modulates social dominance (33) and maternal separation reduces adult dominance in a competition for water in water-restricted mice (34). Furthermore, chronic restraint decreases social dominance in the tube test (35) and severe immobilisation stress heavily reduces social dominance in anxious mice (36). Another study reported the opposite effect of stress on dominance, with maternal separation resulting in increased social dominance in the tube test in rats (37). Therefore, dominance can both regulate the response to stress but also be modulated by it, which indicates the complexity of the neural circuitry regulating this social behaviour, as well as its reactivity to corticosterone.

### F_1_ male and female offspring do not show changes in social conditioned-place preference

Male and female offspring were tested as juveniles in the social conditioned-place preference, and they displayed expression of social reward, with no differential effects of the paternal corticosterone exposure. Social reward has been proposed to drive the approach towards socially relevant environments, as well as the avoidance from predicted social isolation, and is expressed during youth in social animals (38). Through the acquired social experiences resulting from this behaviour, the social reward would then influence the development of adult sociality (39). Stress can modulate social reward, as it has been shown that foot-shock stress exposure reduces social reward expression (40). Stress exposure has also been found to impair sociability (41), with increased aggression in males and social withdrawal in females (42). The effects of stress on adult sociability could be due to impaired acquisition or expression of social reward, although further studies are needed to investigate this hypothesis. Nevertheless, differences in social reward were not observed in our model, which indicates that the changes in adult social behaviour observed in these mice are not due to altered social developmental trajectories.

### Male F_1_ offspring show lower social dominance and increased attractiveness

Male offspring were tested as adults in the social dominance tube test and mate-choice test. PatCort F_1_ offspring showed reduced social dominance in the tube test, like their fathers. This effect was expected, due to their increased anxiety-like behaviour, as studies have showed that anxiety affects social behaviours (43,44), although other studies did not show association between anxiety-like behaviour and social rank in the tube test (45). These mice also showed increased attractiveness, which was not expected. It is important to consider that the setup we used for the mate-choice test intentionally did not let the males display territoriality, compete over mating opportunities, or initiate an interaction with the female, rendering them passive to female mouse discretion. Therefore, the results in attractiveness could have been different if a competitive setup had been used where the males could display their dominance.

Both MUPs and ultrasonic vocalisations (USVs) have been shown to attract females (46,47). When males are exposed to females, they emit USVs as part of a ‘male song’ (48). Additionally, males excrete MUPs in their urine. These proteins have pockets that bind volatile pheromones, but they also act as non-volatile pheromones. MUP expression is complex and conveys a plethora of information, ranging from sex, health status, individuality, and attractiveness (49). By using different setups in the mate-choice test, we tried to determine the key features underpinning, and potentially driving, the increased attractiveness in the PatCort mice. This may include the USVs being derived from the physical presence of the mice in the apparatus, and the MUPs being derived from the presence of soiled bedding. For instance, in our ‘standard’ setup, where we added both mice and their soiled bedding, we accounted for both USV and MUPs.

USVs have been shown to travel short distances and therefore are more relevant for short-distance communication (50). MUPs and other olfactory-based signals, on the other hand, can travel long distances and have therefore been associated with long-distance communication (46). In a natural setting, pheromone marks scattered around a certain area would indicate to a female mouse an attractive socially relevant stimulus. If there was a male mouse nearby, he would then vocalise once the female mouse approached him. Therefore, it is possible that in the mate-choice test setups we used, olfactory marks devoid of pairing with USVs, or vice-versa, could have not been enough to signal attractiveness, hence no differences were observed in the ‘bedding-only’ and the ‘mouse-only’ setups. Additionally, we assessed the male interaction response towards the approaching female in the ‘standard’ set up, to determine whether a differential interaction could have accounted for the changes in attractiveness. However, no differences were observed.

### F_2_ male and female grand-offspring do not show changes in social conditioned-place preference

Following the same experimental design, male and female grand-offspring were tested as juveniles in social conditioned-place preference, and male offspring were assessed as adults in the social-dominance tube test and mate-choice test. However, despite the previously observed increase in depressive-like behaviour in the adult male mice, no differences were observed for any of the social behaviours tested. This shows the limited heritability of the effects that the paternal corticosterone exposure has on social behaviours, which is also observed in other studies, with phenotypes spanning across only one generation following the environmental exposure (10).

### F_1_ offspring mRNA sequencing does not show overt changes in gene expression

Following the reduced social dominance in the PatCort male offspring, we investigated gene expression in the prefrontal cortex of these mice, to investigate potential transcriptomic regulation underpinning this behaviour. We chose the prefrontal cortex because of the association between this brain region and social dominance. For instance, studies have shown that the synaptic efficacy regulated by AMPA receptors in this region controls the expression of social dominance (51), that the increase in social dominance as assessed in the tube test accompanies modifications of the stable actin fraction in synaptic spikes in this region (37), and that social dominance is followed by differential gene expression in this region (45). Lastly, neuronal population activity in the mPFC predicts social rank and success in competitive settings (33).

However, no differentially expressed mRNAs were detected after a rigorous False Discovery Rate correction for multiple comparisons. This effect might be due to four possibilities: 1) Animals were culled at baseline (without acute stress). Unpublished data from our group showed that PatCort males do not differ from controls in baseline plasma corticosterone, but only after a restraint stress. 2) Sequencing was done from bulk-tissue RNA. When considering the heterogeneity of the cellular populations in the prefrontal cortex, it is possible that cell-specific differences in gene expression are not detected (i.e. ‘diluted out’) by bulk-mRNA sequencing. 3) Differential gene expression occurs during development, and we only assessed adult PFC. 4) We only assessed adult PFC, and transcriptomic changes may have been present in one or more other relevant brain areas. We hypothesise that these mice exhibit behavioural changes in adulthood due to differential developmental trajectories that result in subtle neuronal changes, such as modifications in synaptic architecture, spine density or dendritic arborisation. Therefore, differences in gene expression may have been present during critical periods of development, such as during embryonic, early postnatal and/or adolescent stages.

### F_1_ male offspring MUP protein analyses

Urine was collected from adult male F_1_ offspring for quantification of MUP proteins. PatCort mice showed reduced levels of specific MUP bands observed after separation in the SDS-PAGE. Three different bands with molecular weight of around ∼17 kDa to ∼23 kDa were identified, which is similar result to what have been described before (18,20). The band with the lowest molecular weight has been shown to be present in males only and to be the most relevant to signal male attractiveness, and it has been named ‘Darcin’ (18). In our dataset we have observed a reduction in the Darcin and the small band MUP populations (the band with higher molecular weight).

Importantly, contrary to the current literature (18,52), in the mate-choice test we did not observe a correlation between urinary Darcin intensity and male attractiveness. Mate choice is a complex decision that depends on the integration of multiple sensory, affective and cognitive systems (46). Therefore, we hypothesise that although PatCort males exhibit lower levels of Darcin in their urine, other factors might be modulating their attractiveness. Indeed, the different setups we used for the mate-choice test indicate that a combination of both olfactory and auditory stimuli seems to be necessary to signal the differential attractiveness in these mice. Additionally, the highly individual and complex urinary MUP pattern, which is pronounced in wild mice, indicates that female mice use this information for additional profiling of the male quality (53), and therefore could modulate the attractiveness signalled by Darcin alone. It is also important to consider the role of volatile pheromones that bind to and are slowly-released from MUPs, which can have their concentration proportionally altered by social status (54). Due to the reduced social dominance in the PatCort mice, it would be interesting to profile the volatile pheromone content in their urine.

### F_1_ male offspring *Mup* mRNA expression analysis and correlation with MUP bands in SDS-PAGE

Due to the changes in MUP protein expression, we collected the liver from the adult male offspring for *Mup* mRNA expression assessment. No differences were observed between the groups for *Mup3*, *Mup20*, class-B *Mups* and *Zhx2*. However, there was a very strong correlation between *Mup3* gene expression and urinary MUP output in the PatCort only, with no correlation in the PatCT.

The MUP protein concentration normalised by creatinine output, as analysed in this study, is determined by a range of factors. To begin with, the expression of *Mup* genes is induced by many different factors, such as testosterone (55), pulsatile growth hormone (21), and circadian glucocorticoid (56). Social factors also modify the expression of MUPs, with social dominance proportionally affecting MUP production (57), possibly due to changes in testosterone levels (55). The differential transcription of *Mup* gene paralogs results in the large diversity of MUP proteins observed in the urine (58), and one of their known transcription factors is *Zhx2* (22). There is no evidence of post-transcriptional processes regulating MUP protein concentration, as the mRNA expression predicts the urinary protein output (58), and MUPs are not reabsorbed in the kidney (59). Lastly, creatinine levels relate to the muscle mass and are a marker of glomerular filtration (55), and were used in this study to normalise the absolute MUP protein levels in the urine. This normalisation accounts for urine dilution, which can also be changed by social status, with submissive mice reducing their urine production and subsequently increasing their urinary creatinine concentration (57). Therefore, the urinary MUP concentration normalised by creatinine output represents the instantaneous MUP expression relative to the protein levels in the body.

The very strong correlation between *Mup* gene expression and protein output in the PatCort indicates that the overall effect of the factors described above should be homogeneous across this group, resulting in a deterministic association between *Mup* gene expression and its protein output. However, despite no differences in gene expression between the groups, MUP protein is lower in the PatCort. More studies would be necessary to determine the regulatory mechanisms underpinning this result. It is interesting as well that most measures of MUP band levels and *Mup* gene expression have higher variability in the PatCort group only, compared to the PatCT counterpart, which suggests that paternal-Cort exposure affects the expression of these genes, but not homogenously.

Additionally, the primers we used for detecting class-B *Mups* aligned with a range of different *Mup* genes (*Mups 1*, *2*, *7*, *8*, *9*, *10*, *11*, *12*, *13*, *14*, *15*, *17*, *18*, *19* and *22*) to cover some of the *Mup* genes that are evolutionary related (60). Therefore, we may have missed resolving individual *Mups* that are also implicated as female attractants. However, it is important to notice that Darcin (*Mup20*), *Mup2* and *Mup3* are highly expressed compared to the other *Mup* genes (20).

### No differences in MUPs in male F_2_ grand-offspring

Despite no differences in attractiveness observed in the male grand-offspring, urine was collected from these mice for quantification of MUP proteins. As expected, no differences were observed in MUP protein concentration, and neither for the bands separated in the SDS-PAGE. Whilst we know that F_0_ paternal-corticosterone treatment can have effects that transmit to F_2_ grand-offspring, particularly with respect to depression-like behaviour (13), it appears that this transgenerational epigenetic inheritance is specific and does not generalise to the social behaviours and MUP expression that we now report as changed in the F_1_ offspring.

### No differences in methylation of *Mup* genes in male F_0_ sperm

Sperm DNA methylation was assessed through DNA long-read sequencing to determine whether the increased male offspring attractiveness and reduced urinary MUP protein levels could be due to the inheritance of Cort-treatment-induced altered DNA methylation. Although most of the parental DNA marks get erased during early development due to the embryonic reprogramming (61), it has been suggested that certain DNA marks can escape this process (62), as it has been shown to occur in regulatory regions of glucocorticoid and estrogen receptors (63). Additionally, a previous study have shown increases in the DNA methylation of *Mup* genes in adult mice, induced by early-embryonic manipulations, which also resulted in repression of *Mup* transcription (24). However, no overall differences in CpG methylation spanning the *Mup* gene cluster were found in our study, which suggests that other regulatory mechanisms underlie the decreased MUP protein levels in the PatCort male mice.

### Conclusions

Overall, these new results have implications for our understanding of adaptiveness in the context of the epigenetic inheritance, as social interactions are known to contribute to fitness in mice (64). More specifically, social hierarchy can affect survival due to differential access to resources (65) and modulation of the HPA axis responsiveness (66) and, together with male attractiveness, they can impact access to mating opportunities (67,68). The sons (F_1_ offspring) of corticosterone-treated mice showed reduced dominance and, although they showed increased attractiveness, it was only when they were in proximity to their urinary marks. Therefore, although it is not possible to establish the causality between low urinary output and dominance, we hypothesise that in a natural environment, due to their lower dominance, these mice would have lower total urine production and success in marking territory, which has been shown to affect reproductive success in wild mice (69). These male offspring (whose fathers had elevated stress hormone levels) would also have a decline in their survival rates, with the overall effect of reducing their survival.

Aspects of epigenetic inheritance in mammals are still met with some scepticism, with one of the questions being why such inheritance evolved if its impact is rarely observed across many generations (70). However, the hypothesised decrease in survival proposed above due to the reduced social dominance observed in the PatCort mice suggests that by modulating endophenotypes that determine fitness, epigenetic inheritance could impair reproduction and survival, which could then heavily impact the generations to come, even though the changes in behaviour are observed in only one generation. This ‘trans-populational impact’ has been suggested in mice before (3), and has been shown to occur in *C. elegans* (71).

Another factor that can impact adaptiveness and needs to be considered is the mismatch between the environments experienced by the fathers and the offspring/ grand-offspring. Some studies indicate that the epigenetic inheritance could fit into the mismatch hypothesis of disease (72), which posits that changes in the environment during development induce adaptive changes that can prime the individual for that environment (within genetic constraints). For instance, adult-generated neurons born during chronic social stress are uniquely adapted to respond to subsequent chronic social stress (73). However, when an environmental mismatch happens between the timepoints, it can result in maladaptation and disease. Similar mechanisms might be at play behind how the epigenetic inheritance functions. For instance, it has been suggested that when environmental cues are not reliable predictors of offspring environment, in the case of environmental conditions changing between generations, the epigenetic inheritance could instead reduce fitness (74,75).

Lastly, it has been suggested that male attractiveness could have evolved with the aid of epigenetic mechanisms and female mate choice (76). The evolutionary expansion of mouse *Mup* genes is recent (77), occurring due to multiple duplication events (60) that have led to the emergence of many pseudo-genes (78). This indicates selective pressures shaping scent signals relevant for social communication (79), and this is in accordance with olfactory signals that mediate territorial behaviour and sexual selection being under evolutionary pressure (79). Interestingly, one of the first reports on epigenetic inheritance in mice showed differential methylation of *Mup* genes (25), and it has been shown that different sociality levels can have transgenerational effects on MUP expression (20). These studies indicate that one of the avenues to understand the evolutionary relevance of the epigenetic inheritance could be through investigating its regulation of genes that are relevant for sexual selection. For instance, epigenetic reprogramming of reproductive function, driven by early-environment, has been shown in humans (80).

In conclusion, in this study we showed that epigenetic inheritance can modulate social behaviours that are important for determining reproductive success, thus potentially impacting many generations. The present findings, along with other studies investigating epigenetic inheritance, help shed light on the evolutionary impact that this type of inheritance confers. The observed phenotype does not necessarily accompany altered gene transcription in adult offspring (in the limited tissues examined at a single adult stage), which suggests developmental processes are at play. Therefore, it is essential that studies on gene expression focus on the critical developmental timepoints and explore additional relevant tissues and cell population in offspring. Lastly, epigenetic mechanisms underpinning this type of inheritance need to be investigated further, including sperm epigenetics and post-conceptual transfer of epigenetic information via developmental algorithms.

## Methods

### Mice and husbandry

Unless indicated, all mice were housed in groups of 3 to 5 in open-top cages with Aspen shaving bedding (Romania) and 2 sheets of paper tissue for nesting. Cages were changed weekly, and food and water were provided *ad libitum*. 7-week-old *naïve* male and female C57BL/6 breeders were obtained from the Animal Resources Centre (Murdoch, WA, Australia). One week later, male breeders were single-housed and randomly assigned to the control (CT) or corticosterone (Cort) group (total liquid consumption can be found on Supplementary Figure S1). One week before the end of the corticosterone treatment described below, male breeders were tested for the mate-choice and social-dominance tube tests at PND 77. After the designated corticosterone treatment period, CT and Cort male breeders were individually and randomly assigned to *naïve* female breeders and paired for 5 days, after which the males were culled. The females were left single-housed and undisturbed for 19 days, after which they were checked daily for newborn litters. Pups from CT or Cort fathers (Paternal CT – PatCT or Paternal Cort – PatCort groups) were weighed on post-natal day (PND) 7, 14 and 21, during which the boxes were changed. Pups were weaned at PND 28 and were group-housed according to sex and to paternal treatment, with pups from multiple different litters being housed together to avoid litter effects. Male and female offspring were tested for the social conditioned-place preference test at PND 35, and the male offspring were tested for the mate-choice and social-dominance tube tests at PND 77. When the behavioural testing was complete, the male offspring were single-housed and paired with *naïve* females to generate the grand-offspring: grand-paternal CT (GPCT) and grand-paternal Cort (GPCort); the procedure was the same as described above (Supplementary Figure S2). All procedures were approved by the Florey Institute of Neuroscience and Mental Health Animal Ethics Committees (AEC), complying with the Australian Code for the Responsible Conduct of Research and the Australian Code for the Care and Use of Animals for Scientific Purposes. For each generation at least 16 mating pairs were used per group, generating at least 9 litters per group. The n size per group for the behavioural experiments was 16-29. The numbers can be found in Table 1.

### Corticosterone treatment

Corticosterone treatment was as per our published protocols (13,14). Briefly, the Cort group of male mice was given 25 µg/mL corticosterone hemisuccinate (Steraloids Inc., Newport, RI, USA) in their drinking water, changed twice a week, for 4 weeks. Control (CT) male mice received the same drinking water, without corticosterone added.

### Behavioural experiments

#### Social conditioned-place preference

the protocol was adapted from Dölen et al., 2013; Nardou et al., 2019. This test was used to assess social reward, which is the result of the interaction between the approach towards socially salient stimuli, and the avoidance of cues that predict social isolation, which is more easily observed in juvenile mice when social interactions are not affected by sex-specific interests (38). Male and female offspring and grand-offspring, from paternal and grand-paternal CT and Cort treatment respectively, were tested on PND 35. In a locomotor chamber (ENV-510, Med Associates, Fairfax, VT, USA) divided in two similar halves with a door connecting the two halves, two different types of bedding were used to cover the floor in each half (CornCob or Alpha Dri), around 1 cm height. Mice were first assessed in the locomotor chamber and had their activity recorded for 30 min, and the time spent on each side was measured (pre-conditioning). Soon after the exploration, all mice from the same cage were housed together in a new cage with one of the bedding types for 24h, after which they were single housed with the other bedding type for 24h. Mice were then assessed in the locomotor chamber once again (post-conditioning). The time and ratio of the time spent in the social-conditioned bedding during post- and pre-conditioning were evaluated.

#### Mate-choice test

the protocol was adapted from Hoffmann et al., 2020; Mitra and Sapolsky, 2012. This test was used to assess male quality or “attractiveness” as perceived by a fertile female, which predicts the likelihood of first copulation and mating duration (68). Male quality is determined by scent marking (84) and vocalisations (85). Using a 3-chamber interaction test, a fertile female (assessed daily before the test by vaginal smear) explored the apparatus for 10 min. Then, one male from each experimental group was put inside a small cage on each side of the apparatus, and the female explored the apparatus again for 10 min, during which the time she spent interacting with each male mouse was measured. Different set ups were used for this test in order to investigate the underlying factor for an eventual change in attractiveness. The ‘standard’ set up consisted of presenting mice and the respective soiled bedding from their home cage (Figure 1B). The ‘bedding only’ set up consisted of presenting the soiled bedding from their home cage, to assess if pheromones alone determine attractiveness (Figure 1C), whereas the ‘mouse only’ set up consisted of presenting the mice alone, to assess if ultrasound vocalisations or the male interaction *per se* determine attractiveness (Figure 1D). A different female mouse was used for every round of assessment, including for mouse only and bedding only sessions. Females were not tested because this test relies on behavioural responses linked to the development of male secondary sexual characteristics. As follow up on the results found for the F_1_ male offspring, we manually analysed their Mate-Choice trial video recordings to quantify each male’s responsive interaction to the approaching female, to determine if there were differences between the groups for this measure. For this analysis, we quantified the time that each male spent with its snout directed towards the female when she approached the interaction zone.

#### Social-dominance tube test

the protocol was adapted from Tada et al., 2016; Zhou et al., 2016. This test was used to assess social dominance, which underlies agonistic behaviours (87) and can be defined as the capacity to prevail in conflicts encompassing aggression, threats, fights or submission (88,89). The apparatus consisted of a 30-cm long clear plastic tube. During habituation each mouse was trained to go through the tube for 10 times. On the following day, during testing, each mouse from a CT cage faced every mouse from a Cort cage, in a total of 4 interactions per mouse per cage. Each mouse was tested once every after 7 to 9 interactions, so as to allow resting time between the face offs. The number of wins by the Cort group was compared to the null hypothesis on a chi-square test to determine statistical significance. Females were not tested because this test relies on behavioural responses linked to the development of male secondary sexual characteristics.

### Other procedures

#### Urinary component assessments

Immediately prior to being culled, urine was collected from each male by scruffing, and frozen at −80 °C immediately. Mouse urine was thawed and diluted 1/4 in MilliQ water. Previous studies have determined that most of the mouse urinary protein content corresponds to MUP proteins (90). Therefore, MUP concentration was determined using the Quick Start™ Bradford Protein Assay, according to the manufacturer’s instructions. Briefly, diluted urine was incubated with Bradford reagent at room temperature for 5 min, after which it was read at 595 nm in Epoch 2 Microplate Spectrophotometer (Biotek/Agilent). The standard curve was constructed using BSA dilutions ranging from 125 to 1,000 µg/mL (Quick Start Bovine Serum Albumin Standard, Cat. #5000206). Creatine concentration was determined using Creatine (urinary) Colorimetric Assay Kit Cayman Chemical Item No 500701 to account for urine dilution (49), according to the manufacturer’s instructions.

#### SDS-PAGE for MUP protein analysis

The protocol was adapted from Lee et al., 2017; Nelson et al., 2015. Mouse urine was thawed and diluted 1/50 in MilliQ water. Beta-mercaptoethanol and SDS loading buffer were added to each sample and heated for 5 min at 95 °C. 10 µL of each sample was loaded into 4-15% gel and run at 200 V for around 20 min. The bands were stained with 0.1% colloidal blue in ethanol using the Coomassie R-250 staining protocol. A reference comprised of pooled urine from 8 mice was used for semi-quantification, which was run in every gel and used as a normaliser across all gels, after measuring its band intensity with ImageJ (v2.1.0/1.53c).

#### RT-qPCR of *Mup* genes

Mice were killed by cervical dislocation and the right lobe of the liver was dissected, then frozen at −80 °C. Liver RNA was extracted using QIAzol according to the manufacturer’s instructions and quantified in Nanodrop (2000c Thermo Scientific). 1000 ng of RNA was reverse transcribed with SuperScript™ VILO™ cDNA Syntesis Kit (Invitrogen, Cat. #11754050). cDNA was diluted 1/10 for qPCR gene expression studies (Table 4). Relative expression was determined using the comparative ΔΔCt method, with ß-actin as the endogenous control gene. The primers used in this study can be found in Table 4.

#### Offspring prefrontal cortex mRNA sequencing

Mouse prefrontal cortex (bregma +1.42 mm, interaural 5.22 mm) was dissected and snap frozen in dry ice. The RNA was extracted using a standard QIAzol Lysis Reagent (QIAGEN, Cat # 79306) procedure, according to the manufacturer’s instructions. RNA was purified from potential DNA contamination with DNA-*free*™ Kit (Ambion, Cat # AM1906), according to the manufacturer’s instructions. RNA quality was assessed using the Agilent 4200 TapeStation system. Samples with RIN value higher or equal to 7.5 were sent for sequencing at the Australian Genome Research Facility (AGRF) in Parkville, VIC, Australia. Library preparation was performed using Illumina Stranded mRNA protocol, and sequencing was done in the Illumina Novaseq platform on a SP flowcell. 100 bp long reads were single-end sequenced at a depth of 20M to 49M. Adapters were trimmed by the Casava software used by the Illumina platform.

#### mRNA Sequencing data analysis

The Galaxy Australia (v1.0) platform was used for quality control, read alignment and generation of count matrix. Read quality control was done with FastQC (v0.72). Alignment was done with HISAT2 (v2.1.0) (92) using mm10 as the reference. Gene count matrix was generated with HTSeq-count (v0.9.1) (93) with the comprehensive gene annotation of the gencode M25 release (GRCm38.p6) as reference. Lowly expressed genes were filtered out using the default filtering conditions from the edgeR package (v3.34.1) (94,95). Differential analysis expression was done using edgeR, and the volcano plot was made with ggplot2 (v3.3.5) (96). Gene Set Enrichment Analysis was performed using GSEA (v4.2.3) (97,98), with the c2.cp.v7.5.1 gene set database.

#### Sperm collection

For sperm collection, mice were culled by cervical dislocation for immediate dissection of both epididymides. The caudal epididymis was bisected with a clean surgical blade, then immersed into 1.0 mL mt-PBS that was pre-warmed to 37°C and incubated at that temperature for at least 30 mins. Sperm counts were determined, and samples were examined for the absence of tissue debris under a light microscope. Samples were then centrifuged at 400g for 15 mins and excess mt-PBS was carefully removed leaving approximately 100 µL as the final volume. Samples were immediately frozen down and stored at −80°C until subsequent use.

#### Long-read sperm DNA sequencing

High molecular-weight sperm DNA collected from F_0_ males was obtained using MagAttract HMW DNA Kit (QIAGEN, Cat # 67563), following a modified version of the manufacturer’s instructions which consisted of replacing the digestion step with Proteinase K with 40 µL of the reducing agent TCEP (Tris(2-carboxyethyl)phosphine hydrochloride, Sigma-Aldrich, St. Louis, Missouri, United States). 5 µg of sperm DNA from 4 CT and 4 Cort F_0_ mice were fragmented by centrifugation for 60 s at 7200 rpm using g-TUBES (Covaris, Woburn, MA, USA) to obtain 8 kb-long fragments. These fragments were prepared for sequencing with the Ligation Sequencing Kit (Cat # SQK-LSK 109) and sequenced on the PromethION and GridION platforms (Oxford Nanopore Technologies, Oxford, UK) with a mean genome coverage of ∼20X, to ensure methylation calling power.

#### Sperm DNA methylation analysis

Samples were base called using guppy (v4.2.2) and megalodon (v2.2.9) (Oxford Nanopore Technologies Ltd.) against the configuration file “res_dna_r941_prom_modbases_5mC_v001.cfg”. NanoStat (99) was used for data inspection and quality control. Fastq files obtained from megalodon were aligned to the mm10 genome using minimap2 (v2.17-r941) (100), and sam files were sorted and transformed into bam files using samtools (v1.10) (101). The tool f5c (v0.5) (102) was used to call CpG methylation per read, as well as to calculate frequencies of methylation per CpG. A methylation matrix corresponding to the location of the *Mup* gene cluster and its flanking genes Slc46a2 and Zfp37 was generated using the coordinates Chr4: 59,904,830-62,212,385, totalling 11,026 CpGs. Differentially methylated regions (DMRs) between CT and Cort were determined with the program DSS (Dispersion shrinkage for sequencing data) (v2.43.2) (103), as previously described for Oxford Nanopore (104,105). Specific methylation at the *Mup20* promoters was determined using genomic coordinates (Table 5) obtained from the UCSC genome browser promoter track (106). Methylation plots were generated using Nanomethviz (107), and density plots, dotplots and boxplots were generated using ggplot2 (v3.3.5) (96). Statistical analysis of methylation frequencies and data visualisation were performed using RStudio (v4.0.5). The code used for this analysis can be found in the GitHub page (https://github.com/Coracollar/mup_methylation).

#### Statistical analysis

Data was tested for normality with D’Agostino-Pearson tests. If data distribution was normal, it was tested with unpaired t-tests and, if variance differed between the groups, t-tests with Welch Correction were used. If data was not normally distributed, Mann-Whitney tests were used. The mate-choice test was analysed by quantifying the total time spent interacting with each male, which was analysed with the Wilcoxon Matched-Pairs Signed-Ranks test, in accordance with Mitra and Sapolsky, 2012. Additionally, the data was also analysed with the chi-square test for relative preference for each male. The social-dominance tube test was analysed using a Chi-square to assess difference from an expected chance of 50:50, in accordance with Zhou et al., 2016. Comparisons of variance for MUP protein and gene expression data were performed with F test. Statistical analysis was performed using GraphPad Prism 9 for MacOS (v9.3.1). Statistical significance was reached when p < 0.05. Graphs are represented as mean ± standard error of the mean (SEM) for normally distributed data, or median ± interquartile range for non-normally distributed data.

## DECLARATIONS

### Ethics approval and consent to participate

All procedures were approved by the Florey Institute of Neuroscience and Mental Health Animal Ethics Committees (AEC) #19-064, complying with the Australian Code for the Responsible Conduct of Research and the Australian Code for the Care and Use of Animals for Scientific Purposes.

### Consent for publication

Not applicable.

### Availability of data and materials

The sequencing datasets generated and/or analysed during the current study have been deposited in the European Nucleotide Archive (ENA) at EMBL-EBI under accession number PRJEB60786 (https://www.ebi.ac.uk/ena/browser/view/PRJEB60786) and PRJEB60812 (https://www.ebi.ac.uk/ena/browser/view/PRJEB60812).

### Competing interests

M.B.C has received support from Oxford Nanopore Technologies (ONT) to present their findings at scientific conferences. However, ONT played no role in study design, execution, analysis or publication. The other authors declare that they have no competing interests.

### Funding

L.B.H. and C.C.F. are supported by the Melbourne Research Scholarship. A.J.H. and T.Y.P are funded by NHMRC project grants and a DHB Foundation (Equity Trustees) Grant to A.J.H, and an NHMRC Investigator grant [APP11968410] to M.B.C. A.J.H. is an NHMRC Principal Research Fellow whose laboratory is also supported by an NHMRC Ideas Grant and an ARC Discovery Project Grant. M.B.C. and T.Y.P. were co-recipients of a University of Melbourne Midcareer seeding grant.

### Authors’ contributions

L.B.H. planned and conducted the experiments, performed the data analysis and wrote the manuscript. E.A.M. assisted in the animal studies. R.V.H performed the sperm collection and DNA extractions. C.C.F. performed the sperm long-read DNA sequencing study. M.B.C. planned and supervised the sperm long-read sequencing study. T.Y.P. conceived the study, designed the study, supervised data collection and analysis. A.J.H. and T.Y.P. reviewed and edited the manuscript.

## Acknowledgements

We thank Brett Purcell for his assistance with behavioural data collection, Craig Thomson and Helen Huckle for providing beddings for the Social Conditioned-Place Preference, Shanshan Li for assistance with MUP protein analysis, AGRF for sequencing the offspring mRNA, the LIEF HPC-GPGPU Facility for long-read sequencing the sperm samples, and Galaxy Australia and Spartan HPC at the University of Melbourne for providing the platform for bioinformatic analyses. This research was supported by The University of Melbourne’s Research Computing Services and the Petascale Campus Initiative. This research was undertaken using the LIEF HPC-GPGPU Facility hosted at the University of Melbourne. This Facility was established with the assistance of LIEF Grant LE170100200.

## Notes

### Summary of Updates

The manuscript has had major revision.

