## Supplemental figures and tables for "Increased paternal corticosterone exposure preconception shifts offspring social behaviours and expression of urinary pheromones"

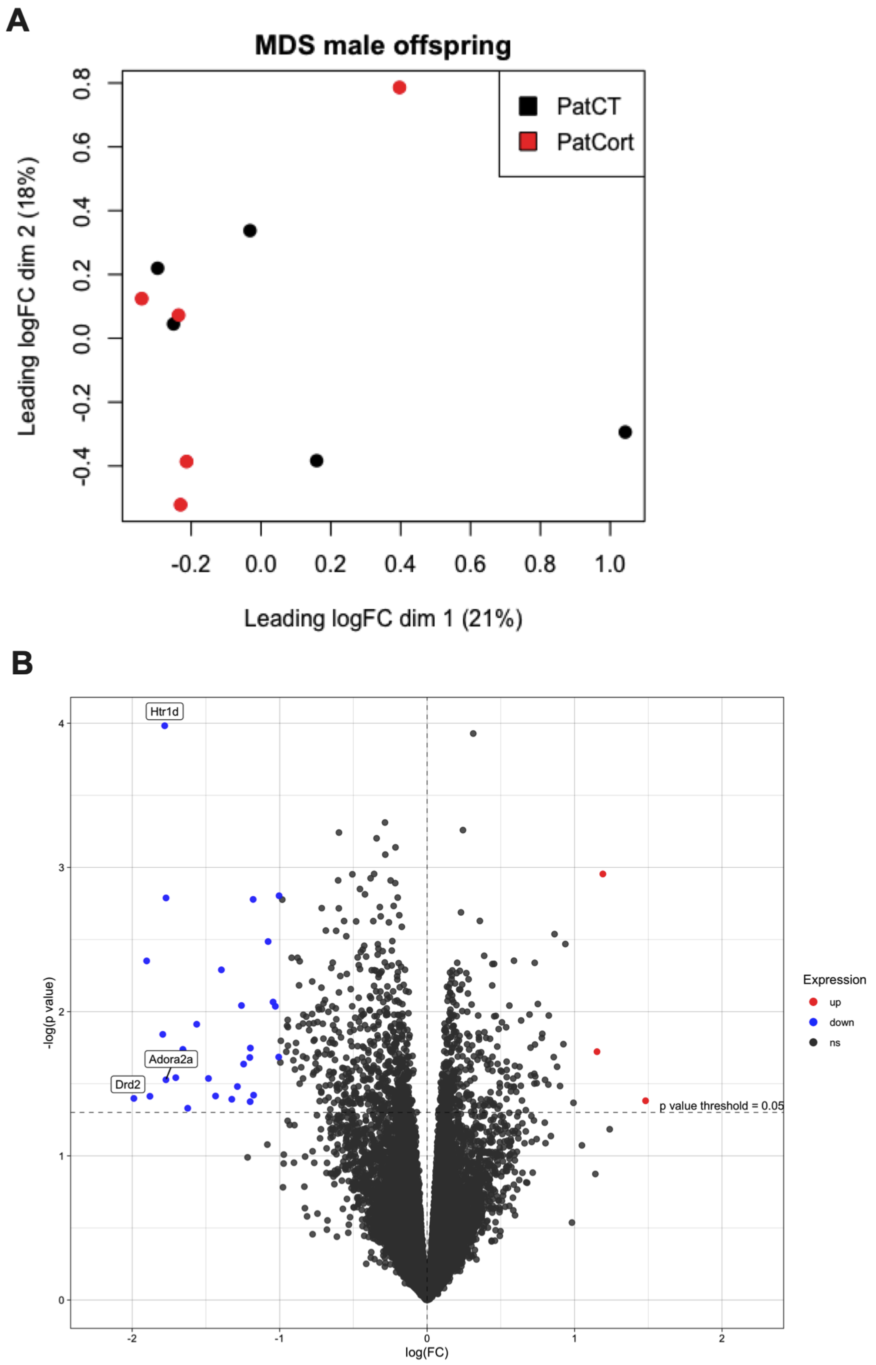


**Figure 7. Assessing effects of paternal Cort-treatment on offspring prefrontal cortex gene expression.** (A), multidimensional scaling plot of distances between the male $F_1$ offspring prefrontal cortex gene expression profiles. (B), volcano plot showing genes with $p$<0.05 and log-fold change threshold of 1 prior to FDR correction. Red: genes upregulated in PatCort. Blue: genes downregulated in PatCort, compared to PatCT.

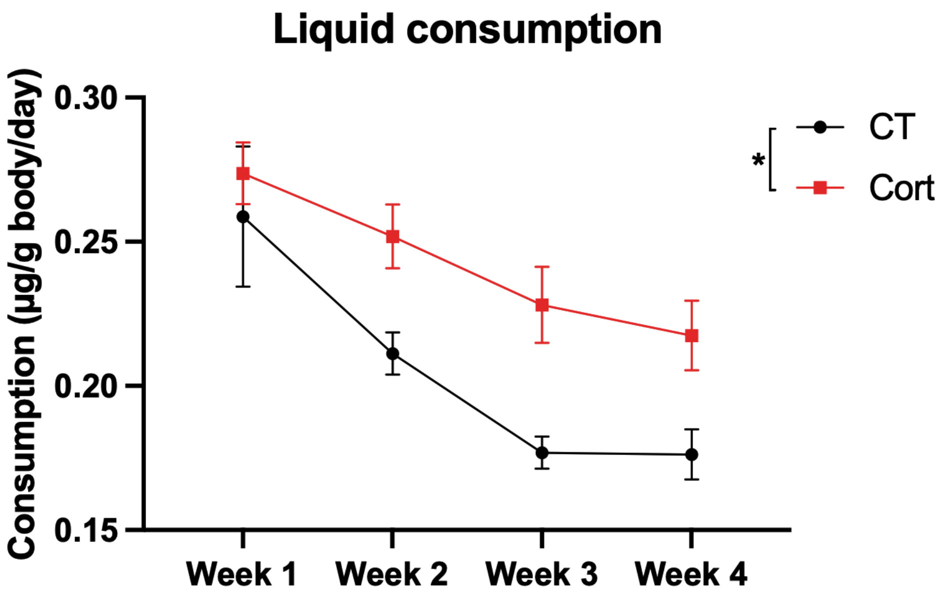


**Supplementary Figure 1. Assessing liquid consumption during Corticosterone treatment.** As expected, Cort-treated mice drink more liquid compared to CT. Repeated-measures ANOVA. CT *n*=16, Cort *n*=16. Group: $F_{(1,30)}$=5.835, *p*=0.04388. * $p$<0.05.

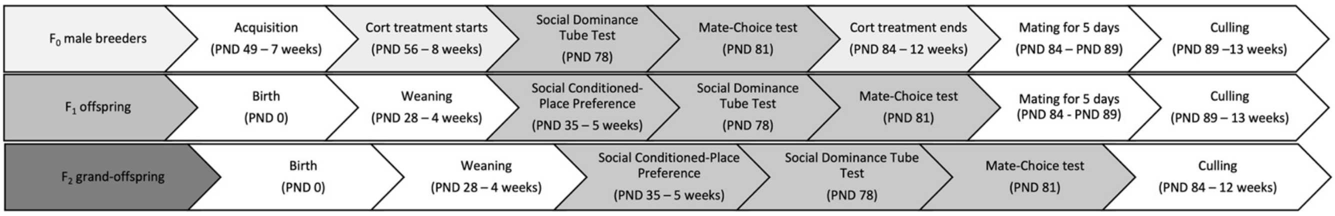


Su**pplementary Figure 2. Experimental design.**

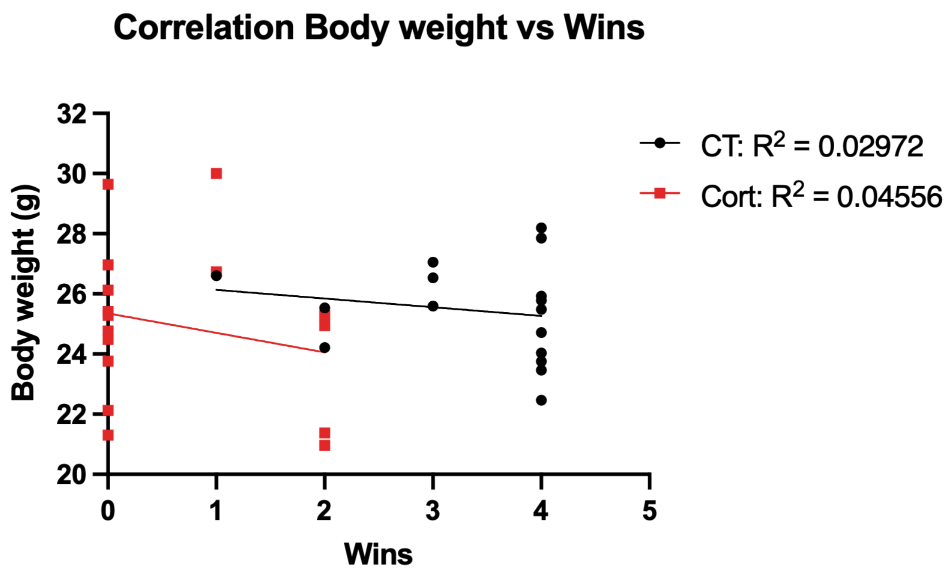


**Supplementary Figure 3. Correlation between body weight and performance in the social dominance tube test..** No statistically significant correlation was found for CT or Cort groups.. Simple linear regression. CT *p*=0.5232, Cort *p*=0.4273.

**Table 1. Number of litters and pups per group.**

| Group | Number of litters | Number of pups |
|---|---|---|
| PatCT | 21 | 162 |
| PatCort | 17 | 125 |
| GPCT | 9 | 66 |
| GPCort | 14 | 92 |

**Table 2. List of genes with $p$<0.05 in the PFC of PatCort male mice prior to FDR correction.**

| gene name | logFC | logCPM | F | PValue | description |
|---|---|---|---|---|---|
| Htr1d | -1.7794088 | 0.1134831 | 26.9624767 | 0.00010408 | 5-hydroxytryptamine (serotonin) receptor 1D |
| Gm16253 | 1.19174884 | -0.6984645 | 16.0603361 | 0.00111027 | predicted gene 16253 |
| 6530403H02Rik | -1.0039713 | -0.4233215 | 14.7331383 | 0.00157162 | RIKEN cDNA 6530403H02 gene |
| Ndst4 | -1.7705306 | 3.04068684 | 14.6040417 | 0.0016271 | N-deacetylase/N-sulfotransferase (heparin glucosaminyl) 4 |
| Gm13680 | -1.1797611 | -0.839943 | 14.5172903 | 0.00166564 | predicted gene 13680 |
| Gm43823 | -1.0783109 | -0.9661731 | 12.1374059 | 0.00326629 | predicted gene 43823 |
| Igfbpl1 | -1.9017291 | 0.15683826 | 11.1175445 | 0.00444885 | insulin-like growth factor binding protein-like 1 |
| Nmbr | -1.395177 | 1.13344848 | 10.661352 | 0.00513102 | neuromedin B receptor |
| Gm16299 | -1.0445865 | -0.3589326 | 9.09198402 | 0.00857875 | predicted gene 16299 |
| Gm6473 | -1.2588983 | 1.69703496 | 8.92938195 | 0.00906811 | predicted gene 6473 |
| Map3k7cl | -1.0292819 | -0.8525231 | 8.89272178 | 0.0091828 | Map3k7 C-terminal like |
| Ano2 | -1.5624926 | -0.210955 | 8.07490909 | 0.01222935 | anoctamin 2 |
| Sh3rf2 | -1.7925705 | 0.40555043 | 7.62873312 | 0.01437308 | SH3 domain containing ring finger 2 |
| Dlk1 | -1.1985842 | 0.55636387 | 7.04088197 | 0.01788999 | delta like non-canonical Notch ligand 1 |
| Gm17794 | -1.6562058 | 0.20614033 | 6.98415481 | 0.0182789 | predicted gene, 17794 |
| Ptprv | -1.6538717 | 0.15827073 | 6.93394202 | 0.01863129 | protein tyrosine phosphatase, receptor type, V |
| Tspan8 | 1.15212195 | -0.8936078 | 6.88796378 | 0.01896083 | tetraspanin 8 |
| Gm5829 | -1.0064098 | -0.6207619 | 6.66503891 | 0.02065689 | predicted gene 5829 |
| Rps15a-ps6 | -1.202439 | -0.9658333 | 6.64615544 | 0.02080842 | ribosomal protein S15A, pseudogene 6 |
| Draxin | -1.2443218 | -0.5397824 | 6.37744503 | 0.0231097 | dorsal inhibitory axon guidance protein |
| Syndig1l | -1.7044298 | 2.80947959 | 5.8359385 | 0.02870111 | synapse differentiation inducing 1 like |
| Glp1r | -1.4826886 | 0.49739511 | 5.80394371 | 0.02907767 | glucagon-like peptide 1 receptor |
| Adora2a | -1.7710575 | 2.99674199 | 5.75323668 | 0.02968623 | adenosine A2a receptor |
| Lrrc10b | -1.2865551 | 2.85216404 | 5.49122498 | 0.03307592 | leucine rich repeat containing 10B |
| Slc35d3 | -1.176682 | 0.80899848 | 5.16393751 | 0.03796042 | solute carrier family 35, member D3 |
| Gpr6 | -1.434612 | 1.18472521 | 5.13221195 | 0.03847698 | G protein-coupled receptor 6 |
| Gm47283 | -1.8795921 | 1.28645542 | 5.12284307 | 0.03863109 | predicted gene, 47283 |
| Drd2 | -1.9885276 | 2.27041218 | 5.04282907 | 0.03997688 | dopamine receptor D2 |
| Cd4 | -1.324498 | 1.13303266 | 5.01099357 | 0.04052746 | CD4 antigen |
| Gm24245 | 1.4814654 | -0.8349585 | 4.95531498 | 0.04151175 | predicted gene, 24245 |
| Gm7908 | -1.199978 | -0.2670917 | 4.92233475 | 0.0421079 | predicted gene 7908 |
| Dnah11 | -1.6232851 | -0.4577555 | 4.68139466 | 0.04677925 | dynein, axonemal, heavy chain 11 |

Legend: genes included with $p < 0.05$ and LFC threshold = 1.

**Table 3. Gene sets detected through GSEA analysis of genes detected in the PFC.**

| NAME | SIZE | ES | NES | NOM p-val | FWER p-val | RANK AT MAX | LEADING EDGE |
|---|---|---|---|---|---|---|---|
| WP_PEPTIDE_GPCRS | 34 | -0.7602396 | -1.7671843 | 0 | 0.1296 | 1059 | tags=32%, list=6%, signal=34% |
| KEGG_OXIDATIVE_PHOSPHORYLATION | 101 | -0.5982546 | -1.6257622 | 4.26E-04 | 0.958 | 1282 | tags=7%, list=7%, signal=7% |
| WP_OXIDATIVE_PHOSPHORYLATION | 45 | -0.655493 | -1.5960314 | 0.005150617 | 0.9946 | 352 | tags=7%, list=2%, signal=7% |

**Table 4. Primers used for RT-qPCR.**

| Gene | Forward | Reverse |
|---|---|---|
| *Mup3* | 5’-GCTTCTGCTCCTGTGTTTGGA-3’ | 5’-CATCAGAGGCTTCAGCAATAGAA-3’ |
| *Mup20* (Darcin) | 5’-GTGCTGCTGCTGTGTTTGGG-3’ | 5’-TGTCAGTGGCCAGCATAATAGTA-3’ |
| Class B *Mups* | 5’-CAGAAGAAGCTAGTTCTACGGG-3’ | 5’-GAGGCCAGGATAATAGTATGCC-3’ |
| *Zhx2* | 5’-AGGCCGGCCAAGCCTAGACA-3’ | 5’-TGAGGTGGCCCACAGCCACT-3’ |
| *ß-actin* | 5’-TATAAAACCCGGCGGCGCA | 5’-ATGGCTACGTACATGGCTGG-3’ |

**Table 5. Genomic coordinates for the *Mup20* promoters, obtained from the UCSC genome browser promoter track.**

| chrom | chromStart | chromEnd | name | score | strand | thickStart | thickEnd |
|---|---|---|---|---|---|---|---|
| chr4 | 61959968 | 61960028 | Mup20_2 | 900 | - | 61959968 | 61959979 |
| chr4 | 62054106 | 62054166 | Mup20_1 | 900 | - | 62054106 | 62054117 |
